# Laser ablation tomography (LATscan) as a new tool for anatomical studies of woody plants

**DOI:** 10.1101/2022.11.07.515046

**Authors:** Israel L. Cunha Neto, Benjamin Hall, Asheesh Lanba, Joshua Blosenski, Joyce G. Onyenedum

**Affiliations:** School of Integrative Plant Sciences and L.H. Bailey Hortorium, Cornell University, Ithaca, NY 14853, USA; Laser for Innovative Solutions (L4iS), Suite 261, 200 Innovation Boulevard, State College, PA 16803, USA; Department of Engineering, University of Southern Maine, 37 College Ave., Gorham, ME 04038, USA

**Keywords:** cell wall, high throughput imaging, laser ablation tomography, lignin, plant anatomy, plant phenotyping, vines

## Abstract

- Traditionally, botanists study the anatomy of plants by carefully sectioning samples, histological staining to highlight tissues of interests, then imaging slides under light microscopy. This approach generates significant details; however, this traditional workflow is laborious and time consuming, and ultimately yields two-dimensional (2D) images. Laser Ablation Tomography (LATscan) is a high-throughput imaging system that yields hundreds of images per minute. This method has proven useful for studying the structure of delicate plant tissues, however its utility in understanding the structure of tougher woody tissues is underexplored.
- We report LATscan-derived anatomical data from several woody stems (ca. 20 mm) of eight species and compare these results to those obtained through traditional anatomical techniques.
- LATscan successfully allows the description of tissue composition by differentiating cell type, size, and shape, but also permits the recognition of distinct cell wall composition (e.g., lignin, suberin, cellulose) based on differential fluorescent signals on unstained samples.
- LATscan generate high-resolution 2D images and 3D reconstructions of woody plant samples, therefore this new technology is useful for both qualitative and quantitative analyses. This high-throughput imaging technology has the potential to bolster phenotyping of vegetative and reproductive anatomy, wood anatomy, and other biological systems such as plant-pathogen and parasitic plant associations.

## Introduction

Understanding the diversity of plant form and function is the motivating principle in the botanical sciences (Haberlandt, 1914). For centuries, botanists have come to understand the morphology and anatomy of plant organs by carefully sectioning plant material, followed by histological staining, and observation through microscopy (Johansen, 1940; Ruzin, 1999). These classical techniques yield fine details on the tissue and cellular levels; however, each step of the traditional workflow is time-consuming, and ultimately yields two-dimensional (2D) microscopy images (see Supporting information Table S1 for a summary of plant anatomy methods). In compliment, scientists usually need to rely on additional techniques to obtain further information on intracellular features, including cell wall composition, cell content, as well as to obtain three-dimensional (3D) reconstructions. Therefore, to thoroughly understand the anatomy of plant organs a large combination of techniques is necessary.

The typical workflow to study the internal structure of plants––i.e., plant anatomy–– with light microscopy usually follows these steps: (1) fixation of fresh plant samples to preserve the architectural integrity; (2) embedding of fixed material to provide the sample with support during sectioning; (3) staining to highlight anatomical features of interests (e.g., cell wall composition) and to generate contrast between tissue types; (4) sectioning on a microtome; (5) microscopy to observe anatomy and (6) imaging to archive anatomical observations (Johansen, 1940; Ruzin, 1999). This general workflow has been applied with great success (Fig. 1a-d, f). Together, the above workflow takes a minimum of two weeks, and each step has its own nuances and pitfalls that must be modified depending on the tissue. Throughout the years, several modifications have been proposed to improve the typical anatomical workflow. For example, multiple alternative fixatives are presented by Johansen (1940) and Ruzin (1999), and there have been several alterations to embedding and sectioning of complex tissues (Barbosa *et al*., 2010; Mozzi *et al*., 2021; Romanov *et al*., 2021). Hard woody samples pose a particular challenge, as these samples must be softened prior to embedding to ensure ease of sectioning downstream; to soften these samples, researchers boil samples in water or in a softens like hydrofluoric acid (Pace, 2019) or Ethylenediamine diluted in water (Kukachka, 1977; Carlquist, 1982) for variable amounts of time which can extends the entire workflow to as long as one month per sample. In some cases, these softening procedures can lead to altering the structure from its native state, either detaching the bark from the wood or by eliminating cell content such as crystals (Pace, 2019). Alternatively, woody samples can be studied by macroscopic analysis which normally requires only polishing the stem surface (Fig. 1g) or wood blocks, followed by imaging through different systems (Table S1), however this technique losses fine-scale detail on cell wall composition, and does not work for herbaceous samples.

**Fig. 1.**
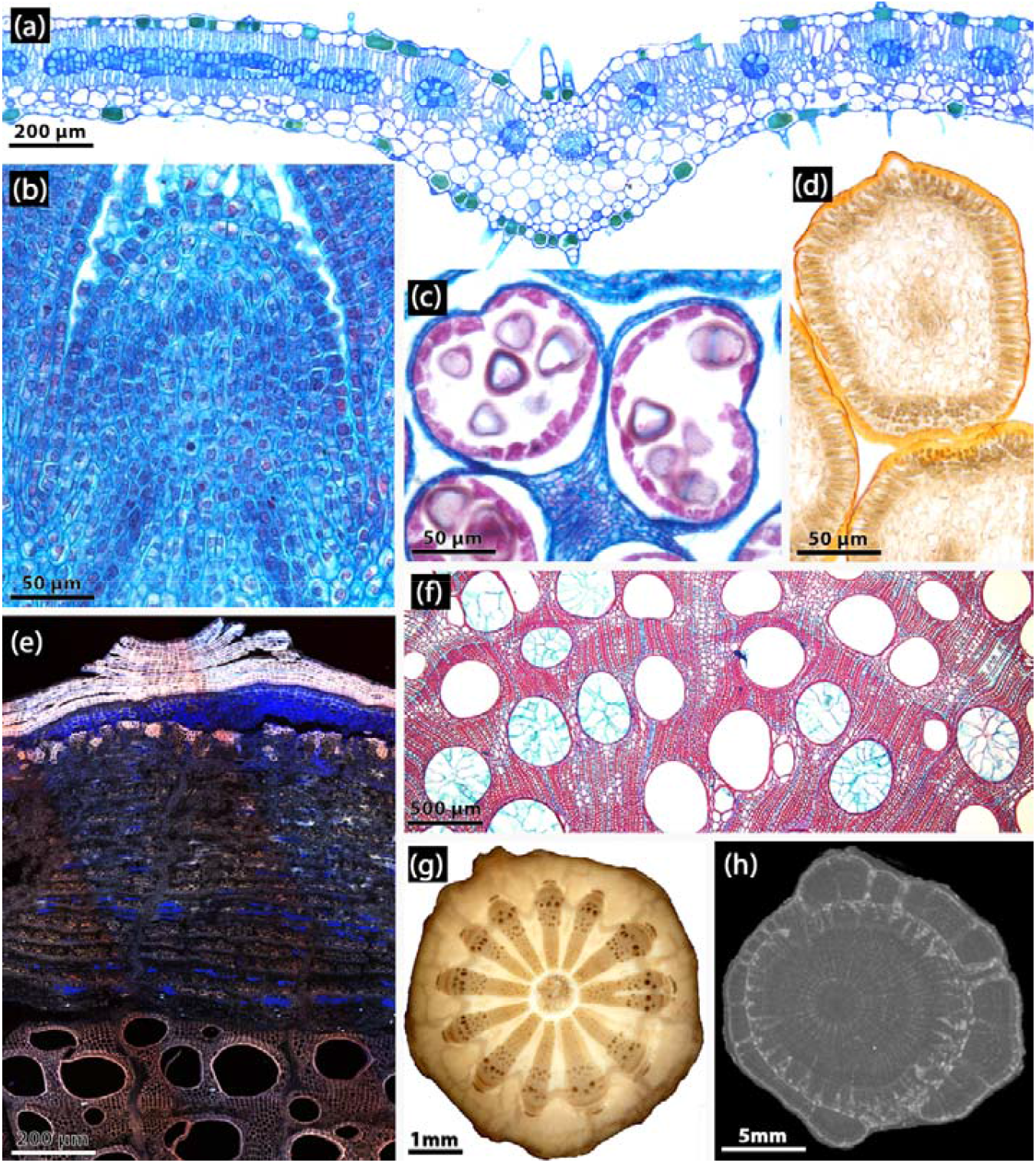
Examples of anatomical images of various plant tissues using different methods applied in plant anatomy. (a-e,g,i) Microscopic analyses. (f) Macroscopy analyses. (h) 3D imaging, X-ray microtomography. (a) *Allionia incarnata* (Nyctaginaceae) – leaf blade processed with historesin embedding, sectioned with rotary microtome, and stained with Toluidine Blue O. (b) *Colignonia glomerata* (Nyctaginaceae) – processed with paraplast embedding, sectioned with rotary microtome, and stained with Safranin O and Astra Blue. (c-d) *Dalechampia alata* (Euphorbiaceae). (c) Detail of anther with pollen grains, processed with paraffin embedding, sectioned with rotary microtome, and stained with Safranin and Astra Blue. (d) Histochemical analysis of secretory glands showing positive result for lipids in the cuticle; sample sectioned with cryomicrotome and submitted to Sudan IV reaction. (e) *Wisteria floribunda* (Fabaceae) – freehand section of the stem, unstained, and imaged with confocal microscopy (maximum intensity projection with three channels). (f) *Dalechampia alata* (Euphorbiaceae), large woody stem processed with polyethylene glycol embedding, sectioned with sliding microtome, and stained with Safranin and Astra Blue. (g) *Menispermum canadense* (Menispermaceae) – polished mature stem, non-stained, processed for macroscopic analysis. (h) *Paullinia micrantha* (Sapindaceae) – stem with successive cambia processed with X-ray microtomography, unstained.

Other methodological approaches that explore anatomical slides in light microscopy is histochemistry, i.e., the application of specific reagents and dyes to detect the main classes of chemical compounds such as lipids and latex (Demarco, 2017; Ribeiro & Leitão, 2020). However, histochemical analysis is preferably applied to fresh tissues, and requires sectioning and staining procedures, as sections should be imaged immediately after test application to avoid altered results. In addition to light microscopy, general anatomy is also frequently explored using wide-field fluorescence or confocal laser scanning microscopy (CLSM) (Fig. 1e) (Prunet *et al*., 2016). Unlike methods applied for light microscope, fluorescence microscopy methods allow analyzing the plant tissues without the need to carry out a staining method. For example, imaging autofluorescence of unstained samples is a simple approach to identify lignified cell walls in woody and non-woody tissues (Kitin *et al*., 2020; Pegg *et al*., 2021; Maceda & Terrazas, 2022), yet the use of fluorescent dyes––e.g., immunohistochemistry with fluorescent secondary antibodies––can reveal the specific localization of other cell wall polymers, including pectins, hemicelluloses, and arabinogalactan proteins (e.g., (Guedes *et al*., 2017).

Within plant anatomy, wood and bark anatomy are particularly laborious because some cells run parallel (axial parenchyma), while others run perpendicular (ray parenchyma) to the plant axis, therefore researchers must section three separate planes i.e., transverse, longitudinal radial and longitudinal tangential to understand the 3D structure of a wood (Brodersen, 2013). Moreover, wood and bark are complex tissues composed of cells with unique contents (e.g., phenolics, starch), and cell wall composition (e.g., cellulose, lignin, suberin), therefore multiple stains must be used to extract these data. Other complex traits that are challenging to understand through traditional anatomical methods are vascular variants i.e., alternative patterns of vascular growth generating odd and intricate morphologies (Bastos *et al*., 2016; Cunha Neto *et al*., 2018; Rizzieri *et al*., 2021), or plant-parasitic interactions where the parasite obtain water and nutrients from the host plant through the complex and dynamic structure, the haustorium (Teixeira-Costa & Ceccantini, 2016; Mylo *et al*., 2021), respectively. To gain a structural understanding of these complex networks, 3D reconstructions have been made using X-ray microtomography (Fig. 1h) and magnetic resonance imaging (Oven et al., 2011; Meixner et al., 2021), however, these methods are time-consuming and result in grayscale data that can miss or conflate different anatomical features. Alternative high throughput methods maximizing the understanding of plant morphology and anatomy continues underexplored in plant sciences.

Laser ablation tomography (LATscan) is a new three-dimensional imaging methodology (Hall *et al*., 2016; Hall & Lanba, 2019). The technology uses a high-powered ultrafast pulsed ultraviolet (UV) laser to remove cross-sections off samples, and an image of the section is captured prior to removal at the laser ablation plane (Fig. 2). A linear stage feeds the sample into the laser ablation plane. The images with UV-induced fluorescence are captured prior to removal. These RGB images allow for easier identification and segmentation of features of interest and are stacked to construct 3D models. LATscan has been used by researchers to study plant and insect anatomy. Yang *et al*. (2019) used LATscan to image maize roots in order to study root phenotypes that improve nitrogen acquisition. The technology was also used to visualize and quantify edaphic organism colonies in maize, barley, bean roots (Strock *et al*., 2019). Morrison et al. (Morrison *et al*., 2020) used LATscan for more accurate measurements of internal wheat grain volumes affected by weevil parasites that grow inside grains. LATscan also helped reveal the structure of specialized storage structures containing symbiotic fungi (e.g., mycangia) in the Ambrosia beetle (Li *et al*., 2018).

**Fig. 2.**
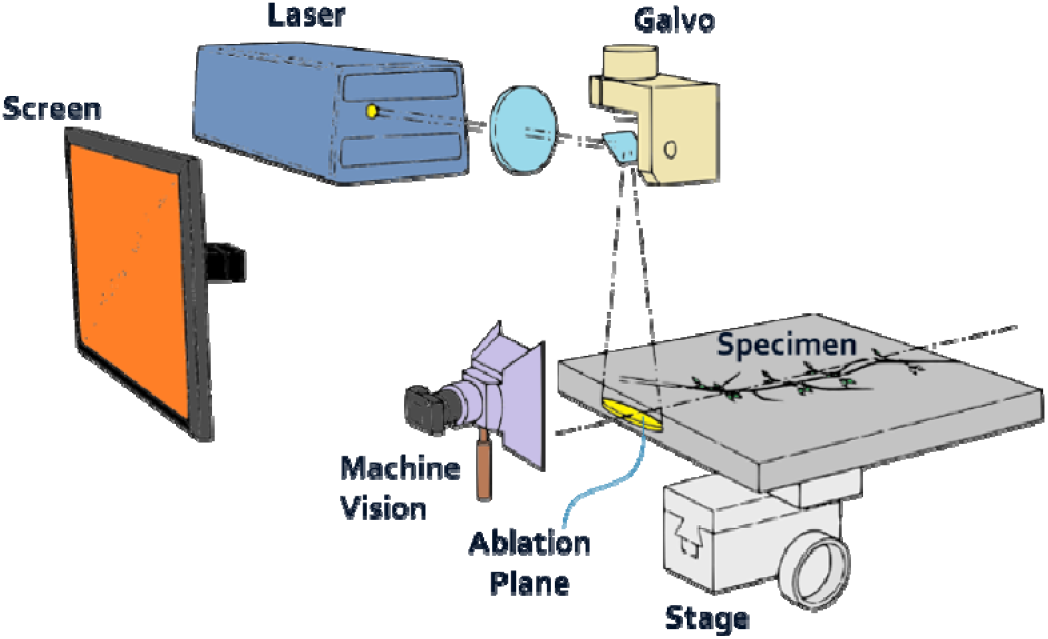
Schematic showing the setup of a LATscan system. The laser beam is guided onto the ablation plane using the galvo, and the specimen is fed in perpendicular to the ablation plane via the stage. As the sample is vaporized in the ablation plane, the remaining section is illuminated via UV-induced fluorescence from the laser, and this is captured by the RGB machine vision system.

Lehnert *et al*. (2022) recently used LATscan to reveal anatomical features in antlion that helped them prove evolutionary adaptation that have made these insects successful in sandy habitats. Schneider *et al*. (2020) used LATscan to quantify root anatomical phenes in their investigation to find the genes that control root plasticity in maize. The technology was also used to quantify and compare the vasculature in oak, beech and spruce branches in order to study hydraulic redistribution under moderate drought conditions (Hafner *et al*., 2017). Therefore, this method has proven useful for studying the structure of small roots, herbaceous plants, and flowers (Martínez-Gómez et al., 2022; Strock et al., 2022), however its utility in understanding the structure of tougher woody tissues is underexplored.

In this study, we demonstrate the potential of LATscan for plant anatomical studies, with emphasis on woody stems of self-supporting and climbing plants (vines). First, we present this technology as a high throughput method to generate 3D reconstructions of woody stems. Next, we explore the potential of LAT as a complementary tool to investigate plant anatomy by comparing the resolution of anatomical data obtained from LATscan to those obtained by conventional methods in plant anatomy. Our results indicate that LATscan is a powerful technology to generate morphological and anatomical features, while revealing differential fluorescent signals corresponding to cellulosic, lignified, and suberized cell walls.

## Material and Methods

### Plant material and study design

Woody stems with secondary growth were obtained from trees, shrubs, and woody vines across different lineages of seed plants (Table 1). In the field, all samples were fixed in formaldehyde-acetic acid-alcohol then subsequently stored in 70% ethanol (Johansen, 1940).

**Table 1.**
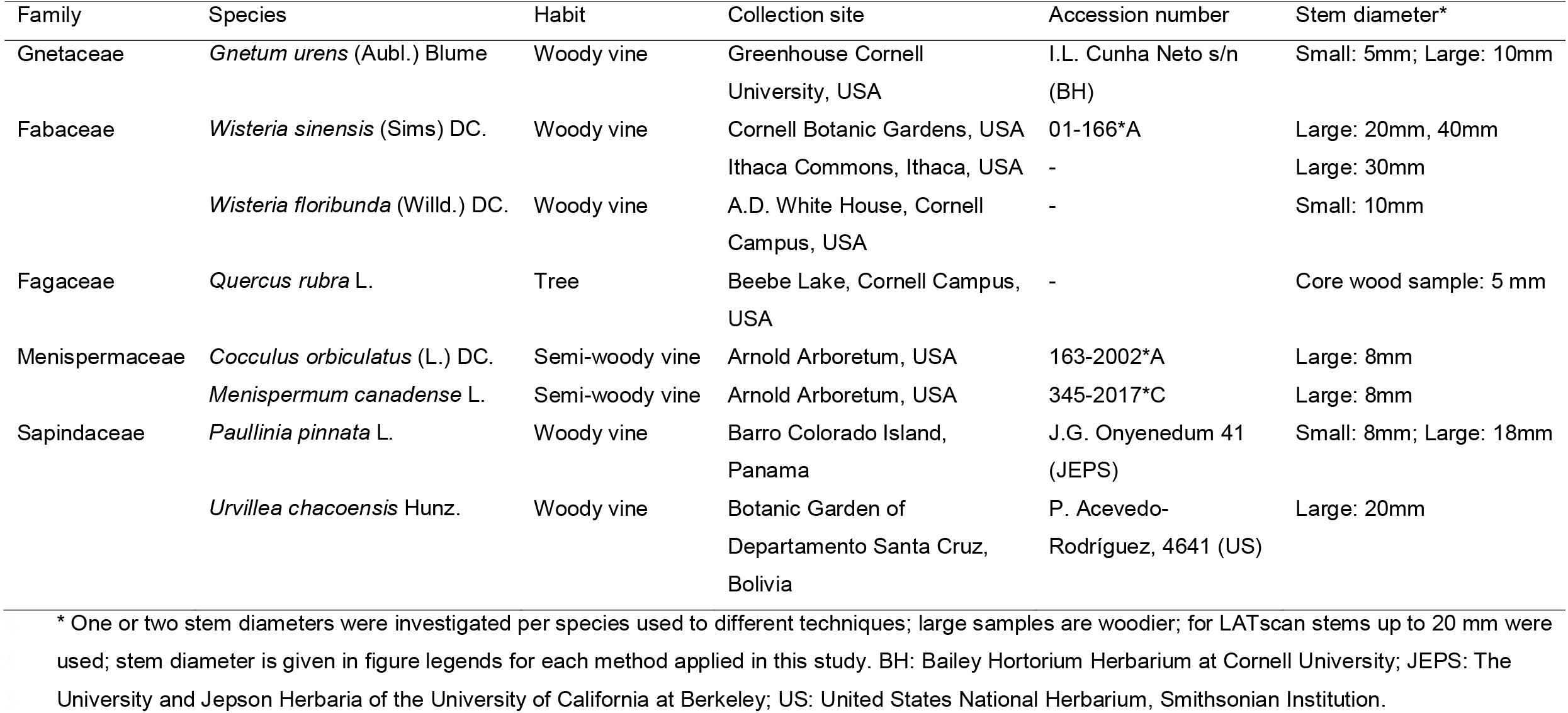
Studied taxa for comparison of traditional anatomical methods and Laser Ablation Tomography (LATscan), including information on site of ollection and stem diameter.

All samples were subject to LATscans and microscopic analyses i.e., generating stained anatomical slides for light microscopy, and unstained slides to detect autofluorescence of cell walls using confocal microscopy. Additionally, all samples were polished and imaged under a stereo microscope for macroscopic analyses of gross anatomy. Below we describe how each of these techniques were optimized for this study.

### LATscan setup and settings (Fig. 2)

LATscan of woody samples were performed at L4iS (State College, PA, USA and Seattle, WA, USA). An ultraviolet (UV) laser outputting at a wavelength of 355 nm was used. The pulse duration of the laser used for the imaging was less than 30 ns and it supplied a pulse energy of approximately 260 μJ. The pulse repetition rate was varied between 15 and 30 kHz. A linear drive stage fed in the sample at increments of 4 μm, and hence the cross-sectional images were separated by that amount. The size of the native RGB images captured was 6720 × 4480 pixels. The magnification resulted in a resolution of 1.1 μm/pixel of the native images.

Image analysis was done at the Lasers and Materials Engineering (LAME) laboratory at the University of Southern Maine (Portland, ME, USA). Image processing was performed using the FIJI software (Schindelin *et al*., 2012). The 3D reconstructions and analysis were performed in FEI Avizo software (v 2019.4 ThermoFisher Scientific, Inc., Waltham, MA, USA).

### Macroscopic analyses

To study the macroscopic characters (gross anatomy and position of important tissues such as periderm, cortex, pericyclic fibers, xylem, and pith) of woody samples, we applied the technique outlined in (Barbosa *et al*., 2021). Briefly, this technique consists of manually polishing the surface of entire or fragments of stems and roots using increasing course grades of waterproof sandpapers (grit grades P600, P1200, P2000), under water. We then imaged polished cross sections using a Nikon SMV1500 stereoscope (Tokyo Japan) with a Nikon Digital Sights Fi-3 camera running Nikon Elements F software (version 4.60).

### Microscopic Analyses

#### 1. Light microscopy analyses

To generate microscopic images, we performed the following steps: (1) freehand sections of each species were generated using a razor blade, (2) sections were stained with Safrablau (see below), (3) sections were mounted with glycerol under a coverslip, and (4) stained slides were imaged using an Olympus BH2 with an Amscope MU1000 digital camera.

#### 2. Confocal fluorescence microscopy

To generate microscopic slides to assess differences in cell wall composition, we performed the following steps: (1) freehand sections of each species were generated using a razor blade (2) sections were either left unstained or stained. The latter sections were stained with either Safrablau or Calcofluor White (see below), (3) sections were mounted with glycerol under a coverslip, and (4) slides were imaged using a IX-83 Spinning Disk Confocal Microscope or (Fig. S1) a Zeiss LSM 710 Confocal Microscope (Figs 1e, 4d,h,l) at the Cornell Institute of Biotechnology’s Imaging Facility.

#### Staining procedures

Safrablau is a combination of two dyes, Safranin-O (a basic or cationic dye with affinity to acid components) and Astra Blue (an acid or anionic dye with affinity to basic components). Anatomical sections double stained with Safrablau (or safranin + astra blue separately) and analyzed under light microscopy, display tissues with red and blue colors, as safranin and astra blue stain mostly lignified/suberized/cutinized or cellulosic cell walls, respectively. Safranin is fluorescent (Excitation= ∼495 nm, Emission =∼ 587nm) while Astra Blue is not. Calcofluor white is fluorescent (Excitation = 365-395 nm, Emission = 420 nm) and labels cellulose which is excited by UV light. Calcofluor white is sensitive to light, thus must be keep in the dark.

To evaluate how different dyes interact with LATscan of plant specimens for anatomical studies we used four staining procedures: 1) unstained samples, 2) Safrablau (9 parts 1% astra blue in 50% ethanol to 1 part 1% safranin in 50% ethanol), 3) Calcofluor White (aqueous, 0.01% w/v, in darkness), 4) Safranin (1% w/v, in ethanol 50%) + 0.01% Calcofluor White. We also tested whether dry or wet samples would respond differently and evaluated different times to stain the samples. In some cases, samples were stained and left to dry before scanning, or they were taken directly from the stain and scanned. Because the whole stem samples are used for LATscan and due to stiffness of most woody stems, some samples were treated longer than usual when compared to the above microscopic anatomical methods. To determine the stains infiltration time, we stained different samples of *Wisteria sinensis*, the stiffest specimen in our sampling. We visually observed the penetration of the dye into the samples, and then analyzed hand sections using confocal microscopy (Supporting information Fig. S1). For blocks 1×1×1 cm wide of *Wisteria*, we soaked samples in Safrablau for one week or more to give enough time for the stain to infiltrate through the whole sample.

Calcofluor White was tested from one hour up to one day. This dye infiltrated faster in samples of similar size stained with Safrablau and were scanned maximum after one day using confocal microscopy. Similar staining procedures were repeated for LATscan. For the combined safranin + calcofluor treatment, we first soaked the samples in safranin, washed in 50% ethanol, and then stained with calcofluor in the dark.

Samples were rinsed at least a few times in distilled water to wash the excess of Calcofluor.

#### Cell wall fluorescence

To quantify the different fluorescence signals of cell walls, we compared cells whose major components of cell walls are lignin or suberin. We compared the chemical composition of cell walls of vessels, G-fibers, and sclerenchyma (lignified pericyclic cells, sclereids and lignified pith cells) to represent lignified tissue, as well as suber and cells in the chemical boundary of the heartwood-sapwood border of *Wisteria sinensis* to represent suberized tissue. We imported images into ImageJ and used the Multi-point Tool to measure the mean fluorescence intensity of 15 individual cell walls representing each of the seven cell types. Data were plotted as a violin plot in R (R Core Development Team., 2021). We tested for differences in emission wavelength means using one-way ANOVA followed by Tukey’s *post hoc* test of honest significant differences (HSD). Values are the mean and *P<* 0.05 was considered statistically significant.

## Results

LATscan is an efficient tool for anatomical studies of woody plants. Our comparisons between unstained and stained samples showed that stained samples with Safranin, Calcofluor or the combination of both dyes did not improve the resolution of images for anatomical investigation (Supporting information Fig. S2). Therefore, below, we describe the application of LATscan for anatomical studies based mostly on unstained samples.

### LATscan yields high-quality 3D reconstructions of complex woody stems under different developmental stages

3D reconstructions of young stems of *Paullinia pinnata*, a woody vine from the maple family, LATscan clearly illustrates the complex anatomy of these stems, which have the formation of multiple vascular cylinders originating in different lobes of the young stem (Fig. 3a-c). LATscan also successfully imaged the complex dynamics of older stems in *P. pinnata*, where the peripheral vascular cylinders undergo anastomoses (Fig. 3d-i) – connections between different portions of the vascular tissues – at the nodal region of the stem (see also Movie S1). 3D reconstructions were also generated for stems of other woody vines with various stem types including stiff samples such as in *Wisteria floribunda*, stems with successive cambia (i.e., multiple increments of secondary xylem and secondary phloem formed in a successive fashion) as in *Gnetum urens*, as well as semi-woody vines with large rays (parenchymatic tissue) such as in *Cocculus orbiculatus* and *Menispermum canadense* (Movie S2).

**Fig. 3.**
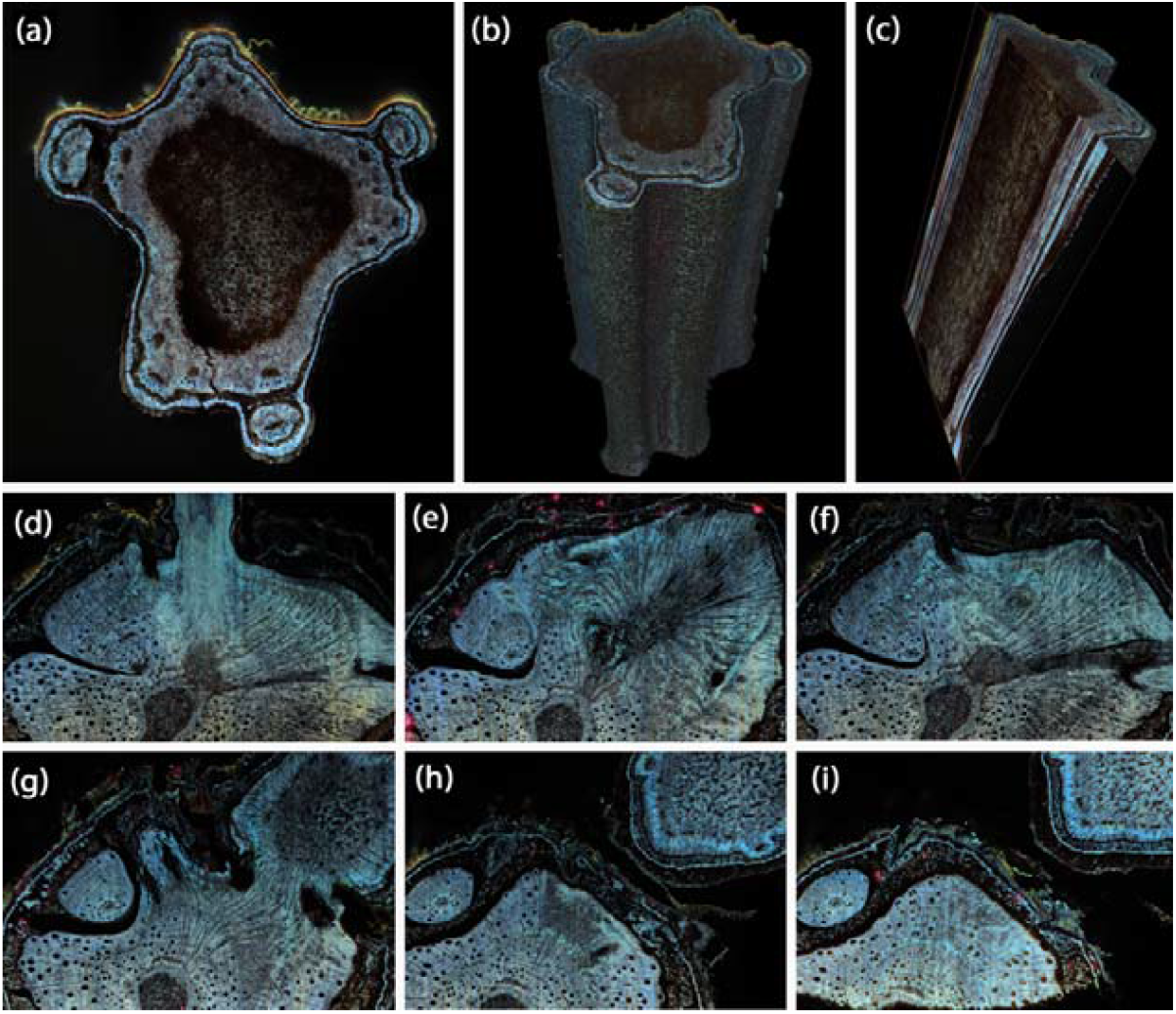
LATscan of the compound stem of the woody vine *Paullinia pinnata* (Sapindaceae). (a-c) Young stem with compound cylinder. Stem diameter: 8 mm. (a) Cross section of the stem in 2D, showing the central vascular cylinder and three peripheral vascular cylinders. (b) 3D reconstruction of the stem. (c) Longitudinal radial view of the stem (computationally sliced), showing mostly pith cells. (d-i) LATscan of the compound stem of the woody vine *Paullinia pinnata* showing anastomoses and splitting between vascular cylinders. Stem diameter: 18 mm.

### LATscan generates substantial structural data for gross stem anatomy descriptions

To assess the potential of LATscan for gross stem anatomy (position of major tissues), we compared images generated through the macroscopic and microscopic methods with LAT images (Fig. 4a-i). From the macroscopic analyses, we observed the main tissues of the stem, including the wood (secondary xylem) and inner bark (secondary phloem), as well as the distribution of vessels, rays, and successive cambia (Fig.4a,e,i). By comparison, LATscan reveals the position of secondary xylem, secondary phloem, and successive cambia like macroscopic images, however they have the additional benefit of more clearly distinguishing the position of the epidermis/periderm, cortex, pericycle, and pith (Fig. 4c,g,k). For example, the macro of *Gnetum urens* is mostly homogenous in color (Fig. 4e), yet the LATscan has high contrast to differentiate this tissue (Fig. 4g). Taken together, these observations indicate that LATscan reveals the same gross stem anatomy features as obtained through the macroscopic method, yet with finer-scale details to differentiate cell and tissue types.

**Fig. 4.**
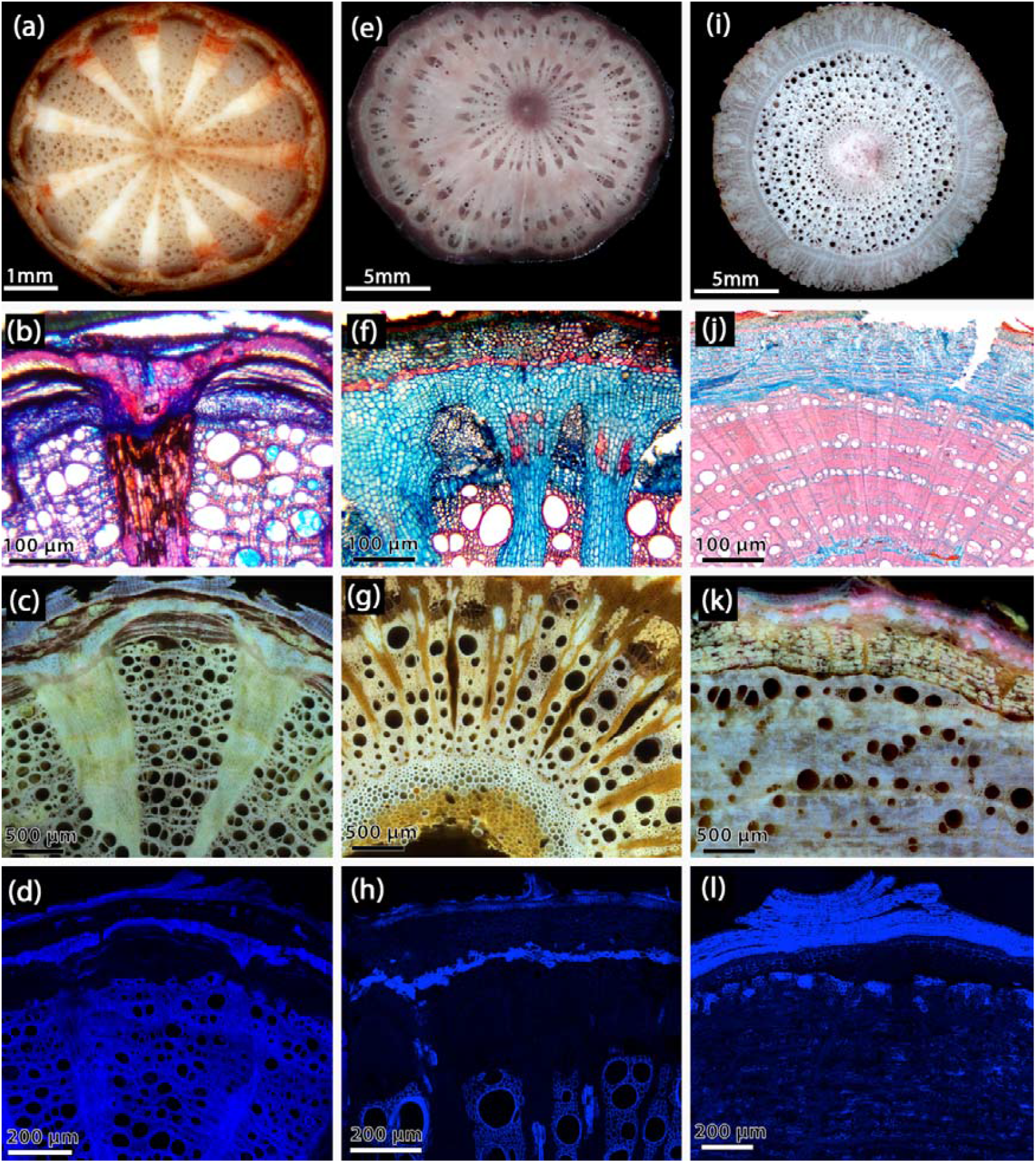
Comparison of cross section images processed for macroscopic analyses, light microscopy and LATscan of woody vines with various anatomical complexity. (a,e,i) Macroscopic images showing mature stems. (b,f,j) Light microscopy, stained with Safrablau. (c,g,k) LATscan of mature stems. Stem diameter: 8 mm, 10 mm, 10 mm, respectively. (d,h,l) Autofluorescence using Laser Confocal Scanning Microscopy (excitation wavelength 405 nm). (a-d) *Cocculus orbiculatus* (Menispermaceae). (e-h) *Gnetum urens* (Gnetaceae). Note G-fibers in the secondary phloem in (f) and (g), and lignified pith cells in (g). (i-l) *Wisteria spp*. (Fabaceae). (i) *Wisteria sinensis* (Ithaca Commons). (j) *Wisteria floribunda*. (k) *Wisteria sinensis* (Cornell Botanic Gardens). (l) *Wisteria floribunda*.

From the light microscopy analyses, we obtained finer details in comparison to macroscopic images, allowing the distinction of cell types (e.g., parenchyma, fibers, vessels) and their shape, size, and distribution of cells (Fig. 4b,f,j). Specifically, light microscopy images revealed, for instance, the phloem cells in *Cocculus orbiculatus* (Fig. 4b) that was identified only as brown patches by macroscopic (Fig. 4a), the presence of gelatinous fibers in the phloem, sclereids in the rays, and fibrous pericycle in *Gnetum urens* (Fig. 4f) and the stratified arrangement of fibers alternating with other cell types in the secondary phloem and fibrous pericycle of *Wisteria floribunda* (Fig. 4j). By comparison, LATscan also reveals these fine details, capable of also differentiating vessels from parenchyma from fibers, with the added benefit of differential autofluorescence (see cell wall composition results below).

LATscan is a powerful tool for secondary xylem and secondary phloem qualitative and quantitative anatomy To investigate the potential of LAT for wood and inner bark anatomy, we compared images generated through light microscopy method to LAT images. We used vines with various degrees of woodiness, diversity stem types (regular growth vs. vascular variants), and periderms.

Microscopic methods are particularly challenging with woody stems, as it requires sectioning plants in three faces (i.e., transverse, longitudinal radial, longitudinal tangential) to understand the 3D structure. However, with LATscan we can generate 3D reconstructions (Fig. 5; Movie S2), and re-oriented them to computationally slice through the sample in either transverse, radial or tangential view to characterize wood and bark anatomy accordingly (Fig. 5, 6).

**Fig. 5.**
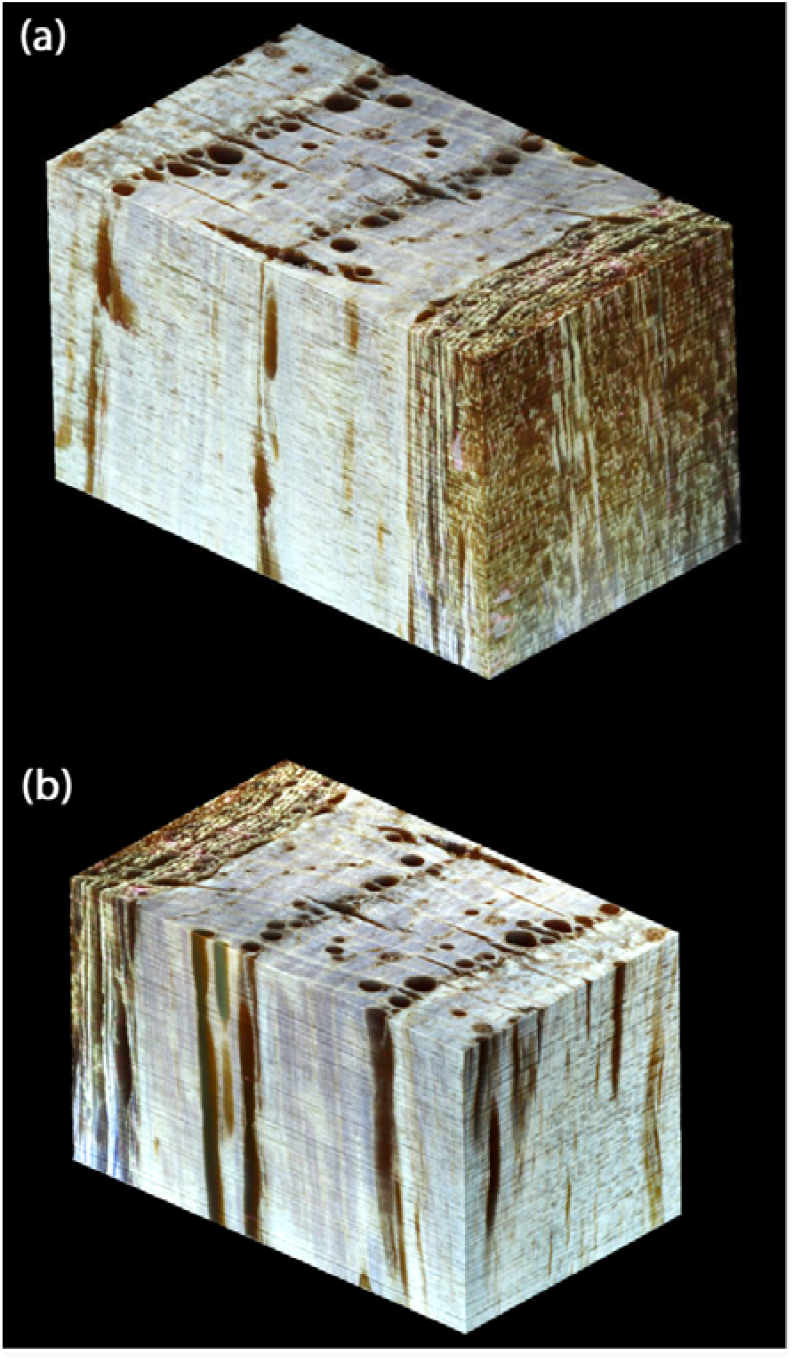
Schematic wood block showing three section planes of *Wisteria sinensis*, reconstructed from LATscan. (a) Top view showing the bark in front view. (a) Top view showing the tangential view of the bark (front view), radial plane (lateral view) and cross view (top). (b) Top view showing the tangential view of the wood (front view), radial plane (lateral view) and cross view (top). Block dimensions: 2.5 × 1.5 × 1.6 mm^3^, with a voxel size of 1 × 1 × 5 micron^3^.

**Fig. 6.**
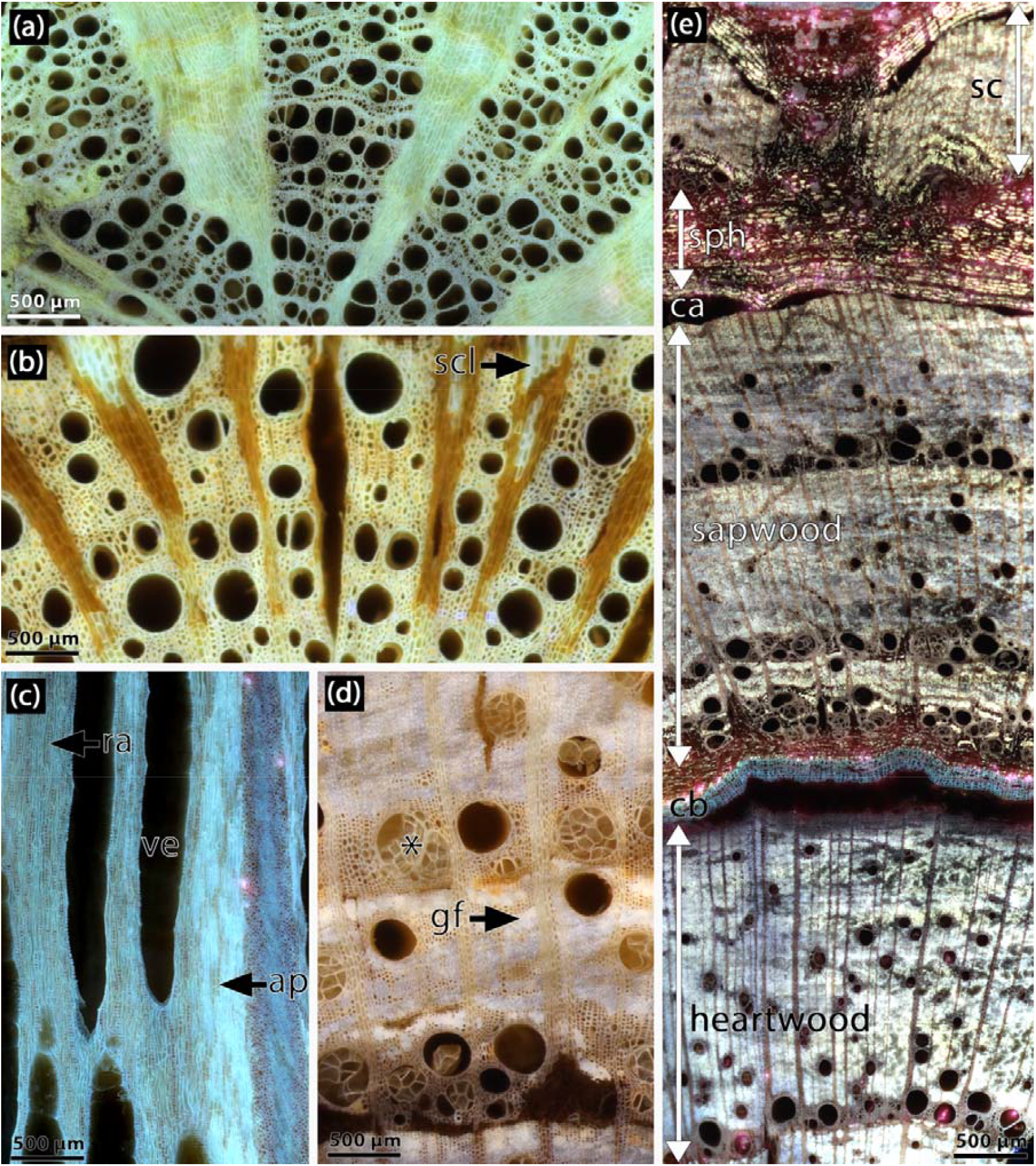
LATscan of woody vines illustrating wood anatomical features. (a) *Cocculus orbiculatus* (Menispermaceae) – wood has indistinct growth rings, wide and narrow vessels solitary or in multiples of two; large rays are formed by parenchymatic cells. (b) *Gnetum urens* (Gnetaceae) – wood with indistinct growth rings, diffuse-porous and solitary vessels; vessels are distributed in a background of fibrous cells; large parenchymatic rays are observed, with groups of lignified parenchyma cells (sclereids). (c) *Quercus rubra* (Fagaceae) – longitudinal tangential section showing short uniseriate rays, axial parenchyma, and vessels. (d-e) *Wisteria sinensis* (Fabaceae). (d) Note ring-porous wood with large vessels produced at the beginning of the growing season, tyloses in large vessels (asterisk), and two types of fibers, the regular fusiform fibers (larger lumen) which intermix with parenchymatic cells, and g-fibers with smaller diameter and a blurry white aspect in this image. (d). General view of a mature stem showing heartwood and chemical boundary (compartmentalization zone with suberized cells), sapwood with growth rings, secondary phloem, and new increments of vascular tissue (successive cambia). ap, axial parenchyma; ca, cambium; cb, chemical barrier (compartmentalization zone); gf, g-fibers; ra, vascular ray; scl, sclerenchyma; sph, secondary phloem; ve, vessel.

Using this method, we can note that in cross section the wood of *Cocculus orbiculatus* (Fig. 6a) and *Gnetum urens* (Fig. 6b) are characterized by relatively large vessels embedded in a background of lignified cells, with large parenchymatic rays.

Sclerenchymatic cells are observed in the rays of *Gnetum urens* (Fig. 6b). *Wisteria sinensis* has growth rings, porous wood (large vessels in the early wood), and a matrix of parenchymatic cells and fibers surrounding the vessels (Fig. 6c-d). Tyloses are common in large vessels of *W. sinensis* (Fig. 6c-d) and the heartwood that appeared as a dark core with macroscopic method (Fig. S3) became anatomically visible with LAT like the rest of the stem (Fig. 6d). Note uniseriate rays and axial parenchyma strands in longitudinal tangential sections of oak (Fig. 6c). We used high-resolution volume renderings to quantify the number and proportion of vessels in *Urvillea chacoensis* (Fig. 7a) using ImageJ (Fig. 7b) and Avizo (Fig. 7c). We found that the cross-section investigated using ImageJ (Fig. 7b) have 55 large vessels (diameter >50 μm), corresponding to nearly 20% of the total cross-section (Fig. 7b). Vessels varied from 53 μm to 230 μm wide (Dataset S1). This type of analysis can be performed for different purposes, as images can be virtually dissected, in any dimension, from the 3D reconstruction (Fig. 7c).

**Fig. 7.**
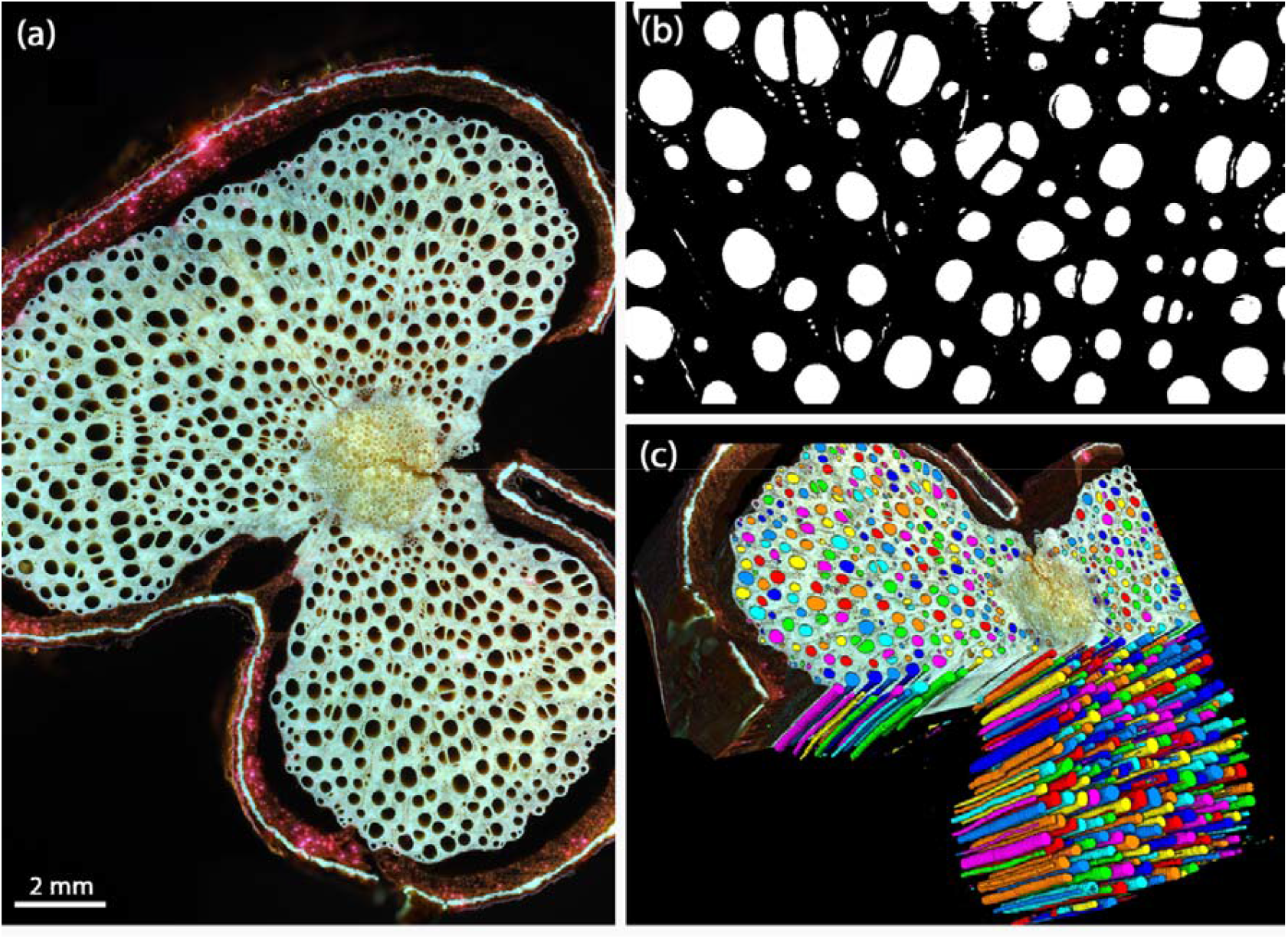
Anatomical images of the stem and wood of *Urvillea chacoensis* (Sapindaceae). (a) Original image generated by LATscan. (b) Binary image after performing threshold analyses using ImageJ, which is the basis for quantifying anatomical features such as vessel diameter. (c) Image after performing quantification analysis using AVIZO.

The bark is divided into inner bark or secondary phloem, and outer bark that includes the pericycle, cortex, epidermis and/or periderm. As for the wood, LATscan revealed different cell types present in these tissues (Fig. 8Aa-e). Conducting cells of the phloem and the sieve-tube shape are still observed along with other parenchymatic cells (Fig. 8a-b,e). Sclerenchyma associated with the phloem is also revealed (Fig. 8a,e), including G-fibers in *Gnetum urens* (Fig. 8b) and *Wisteria sinensis*, where they form seemingly alternate bands with other cell types (Fig. 8c). The pericycle (Fig. 8c), cortex (Fig. 8b,e) and periderm (mostly the suber) are also discernible, especially due to their different fluorescent signals (see results below).

**Fig. 8.**
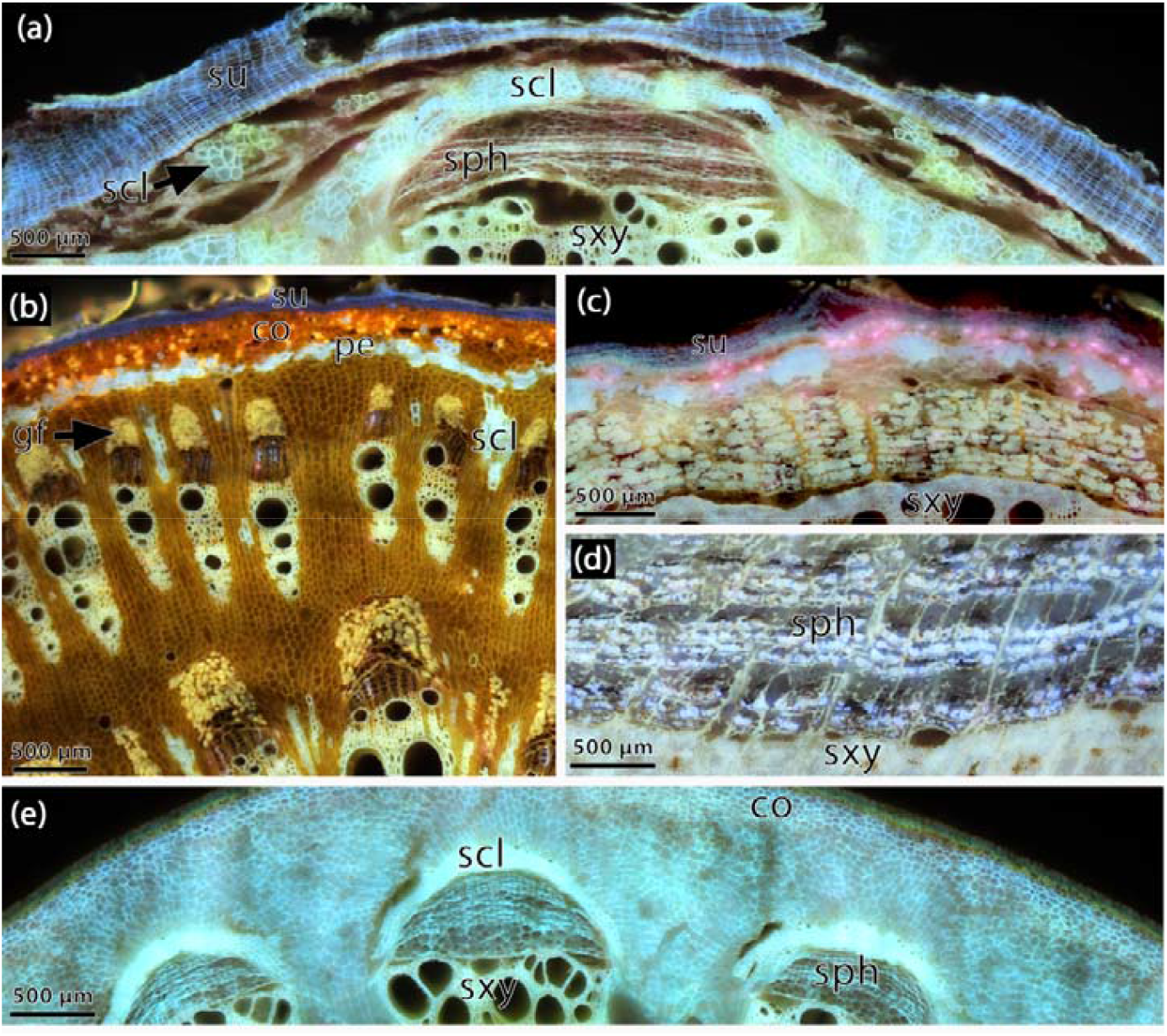
LATscan of woody vines illustrating bark anatomical features. (a) *Cocculus orbiculatus* (Menispermaceae) – secondary phloem formed by thin-walled cells (sieve-tube elements and parenchyma), continuous multiseriate and fibrous pericycle, lignified cluster of cells in the cortex, periderm with several layers of suber. (b) *Gnetum urens* (Gnetaceae) – secondary phloem formed by thin-walled cells (sieve-tube elements and parenchyma) is opposed to conducting cells of the secondary phloem which are separated by large rays; sclereids are present in the rays. Also note continuous fibrous pericycle, cortex and periderm with few layers of suber. (c) *Wisteria sinensis* (Fabaceae) – secondary phloem formed by alternating bands of fibers with sieve-tube elements and parenchyma. (d) *Menispermum canadense* (Menispermaceae) – secondary phloem formed by thin-walled cells (sieve-tube elements and parenchyma) is opposed to conducting cells of the secondary phloem which are separated by large rays. co, cortex; gf, g-fibers; scl, sclerenchyma; sph, secondary phloem; su, suber; sxy, secondary xylem.

### LATscan is a proxy for cell wall composition given their distinct fluorescent signals

To investigate the potential of LAT to reveal cell wall composition, we compared fluorescent signals from confocal autofluorescence microscopy (imaged with 405 nm, 488 nm and 561 nm) of both unstained and stained samples (Fig. 4d, h, l; Fig. S1) to LATscan (Figs 4, 6, 8). Across all species, confocal analysis detects a strong autofluorescence (excitation at 405 nm) of lignified cells (e.g., fibers, vessels, pericycle) and suberized cells (e.g., suber) (Fig. 4d,h,i). This indicates that lignin fluorescence is similar to suberin with intense blue fluorescence under UV excitation (Fig. 4d,h,i; Fig. S1). In general, LATscan displayed different fluorescent signals for cell types with distinct cell wall composition. The differences are here described qualitatively and quantitatively. In unstained samples, xylem fibers and vessels (cells with thick, lignified walls) have a similar signal (Fig. 4c,f,i). G-fibers, which have an additional layer of cellulose that later mature into a lignified layer, have a particular fluorescent signal, which may vary from a golden color in the secondary phloem and cortex of *Gnetum urens* (Fig. 8C), while in *Wisteria sinensis* they display a whitish or bluish color in the wood (Fig. 4k) and bark (Fig. 8d) respectively. Pericyclic fibers (Fig. 4c, 4f; 8a-c), lignified pith cells (Fig. 4i) and sclereids (Fig. 4i, 8c), which are parenchymatic cells that later become lignified, have a similar bright white fluorescent signal across species. Suberin-rich cells in the periderm (Fig. 4c, 4f, 8a, 8c) or in the chemical boundary of the transition from heartwood to sapwood in *Wisteria sinensis* (Fig. 6a) display a blue-ish fluorescent signal. In all species, parenchymatous cells (pith, axial and radial parenchyma, cortex) have a particular fluorescent signal (4c,g,k) differentiating them from sclerenchyma, vessels or periderm.

The fluorescence intensity for the main cell types were determined to investigate the different fluorescent signals. Across species, the analysis showed a mean fluorescence range of 50-129 nm for vessels, 49-181 nm for sclerenchyma, 106-144 nm for G-fibers, 35-158 nm for suber and 74-105 nm for boundary layer in the heartwood-sapwood border (Dataset S2). Among lignified cells, there was a statistically significant difference in mean fluorescence signals between at least three groups (one-way ANOVA, df= 4, *F*= 40.06, *P<0*.*001*). Tukey’s HSD Test for multiple comparisons found that the mean value of suber was significantly different between vessels, G-fibers, and sclerenchyma (*P<0*.*001*) (Fig. 9). There was no statistically significant difference between suber and heartwood (the chemical boundary of *Wisteria sinensis*) (*P*= 0.98), and between heartwood and vessels (*P*= 0.47) (Fig. 9).

**Fig. 9.**
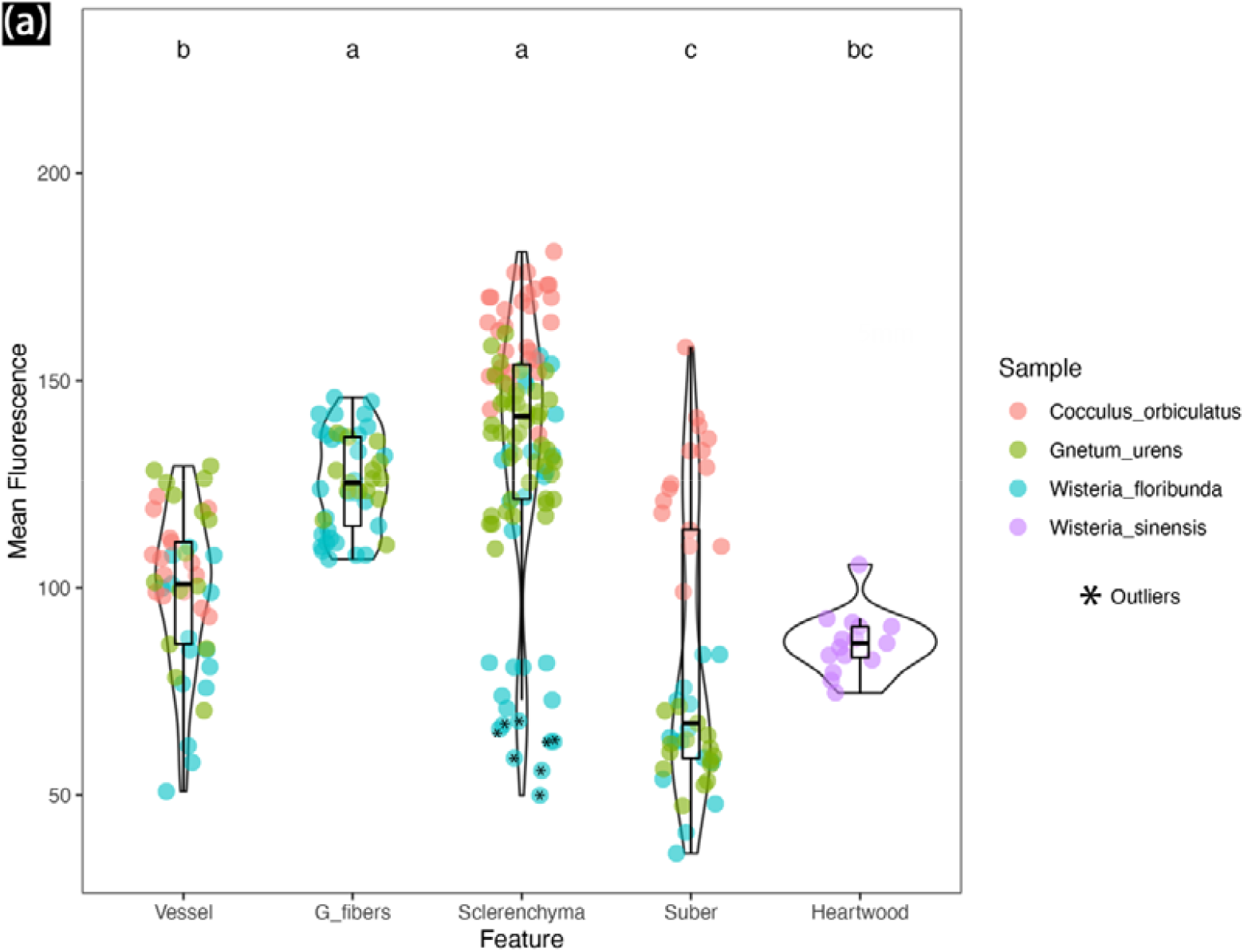
Violin plot of emission wavelength measurements for different cell types analyzed from tree species of woody vines. Different letters indicate significant difference by one-way ANOVA (P<0.05). G-fibers are absent in *Cocculus orbiculatus*. Heartwood was present only in *Wisteria sinensis*.

## Discussion

With the rebirth of comparative plant morphology, there is an increased amount of research focusing on micromorphological studies as plant scientists pursue the question on how organisms evolved and diversified. To understand such morphological complexity of plants, various techniques have been developed or improved to facilitate plant phenotyping (Legland *et al*., 2018; Strock *et al*., 2022). Here, we tested LATscan, a new imaging technology applied in plant sciences, to demonstrate its potential as an alternative tool for qualitative, quantitative, and high-throughput research of woody plants.

### LATscan is a fast and versatile technique for anatomical studies of woody plants

Compared to other techniques, LATscan is an excellent tool for anatomical research for different reasons. First, it is a faster, high throughput technique that produces high-quality 3D reconstructions as well as 2D images suitable for plant anatomical studies. Previous studies demonstrated that LATscan can provide high throughput data of roots, shoots, and inflorescences from herbaceous plants (Strock *et al*., 2019; Martínez-Gómez *et al*., 2022). Here we demonstrated that this technology is also adequate for woody (stiff) samples. LATscan enabled 3D reconstructions of stem samples in different developmental stages, including plants with different degrees of woodiness, habits and from distant lineages of seed plants (gymnosperms and angiosperms). In general, these 3D reconstructions presented higher quality and resolution if compared to other woody stems scanned by X-ray microtomography which can be applied both to dead samples (see Table S1) or in vivo (e.g., to study vessel embolism in woody plants - Brodersen *et al*., 2010; Cochard *et al*., 2015). We believe that in addition to woody samples, LATscan may be suitable for the study of complex systems involving the association of multiple organisms, such as plant-pathogen interactions and the interaction of haustoria of parasitic plants with host plants.

The second reason why LATscan facilitates the study of woody anatomical tissues is because it generates from a single scan information on qualitative, quantitative, and chemical composition of cells and tissues. Qualitatively, the resolution of images allowed cells and tissues to be properly described, similar to the results obtained by images originated from histological slides using light microscopy (See Table S1). We evaluated different species, which are anatomically distinct in terms of distribution and abundance of cell types encompassing stiff and soft tissues (see Fig. 4). This broader sampling permitted to compare diverse wood types, indicating that LATscan presents consistent results independent of wood stiffness. In general, light microscopic images of histological slides yield additional details at the cellular and tissue level that might be unnoticed in macroscopic analysis. Here, we demonstrated that LATscan can generate enhanced data over these two techniques, as was demonstrated for the structural and chemical wall composition of the boundary tissue in the transition of heartwood and sapwood in stems of *Wisteria sinensis*. These cells with unusual arrangement and a blue, fluorescent signal similar to suberized cells in the periphery was interpreted as chemical boundary or barrier zone (compartmentalization system). This phenomenon comprises the formation of structural and chemical barriers in the wood after injuries, wounding, or both, which may function as a protective mechanism against pathogens (Pearce, 1996; Spicer, 2005; Schweingruber, 2007). Our results on fluorescence emission wavelength (see discussion below) corroborates that a major deposition of suberin (instead of lignin) occur in these cells, which is one of the metabolites that may be synthesized by living cells in response to wood injuries (Pearce & Rutherford, 1981; Spicer, 2005). LATscan also proved useful for quantitative wood anatomy, using 2D images for automatic image analysis of wood structure, similar to how histological slides are analyzed (Scholz *et al*., 2013; Ziemińska *et al*., 2015; Von Arx *et al*., 2016).

Lastly, LATscan enabled the identification of cell types with distinct cell wall compositions as a result of contrasting fluorescence signals emitted from these tissues. Our analysis demonstrated that there is a qualitative and quantitative consistency in the pattern of spectral variation, which helped to pinpoint different cell wall components (e.g., lignin, suberin, cellulose). Specifically, statistical analysis helped differentiating suberized and lignified cells across stem types, while mean fluorescence of different types of lignified cells (e.g., pericyclic fibers, sclereids) were not statistically different, which might be explained by the different but continuous levels of lignification in these tissues. We noted that G-fibers which occur in *Wisteria* species (in the wood and bark), and in the bark of *Gnetum urens* had a wide qualitative range in the fluorescent signal, varying from white to blue and gold, but were not statistically different from each other. Biologically, these results may be explained by the different maturation stages of G-fibers, which arise as mainly cellulosic, but can undergo delayed lignification; indeed numerous species have been shown to have lignified G-fibers (Ghislain & Clair, 2017).

The assessment of chemical cell wall composition through direct visualization of spectral bands have been reported using different systems, such as histochemical analysis through autofluorescence using confocal microscopy (Hutzler *et al*., 1998; Donaldson & Williams, 2018) or chemical imaging by confocal Raman microscopy (Gierlinger & Schwanninger, 2006). For instance, Donaldson & Williams (2018) used autofluorescence to characterize the variation in cell wall components including lignin and suberin from healthy, chlorotic, and necrotic pine needles. Similarly, several studies have highlighted the usefulness of cell wall autofluorescence for wood science, because wood cells are naturally fluorescent mostly due to the presence of lignin (Donaldson, 2013; Maceda & Terrazas, 2022). Examples of such studies include the assessment of differences in lignin composition and localization of polysaccharides in the cell wall of normal and compression wood (Donaldson & Knox, 2012; Donaldson & Radotic, 2013) or the topochemical characterization of fibrous, dimorphic and non-fibrous stems of cacti based on lignin composition/ratio (Maceda *et al*., 2019). In general, these studies highlight that fluorescence spectra through confocal microscopy can help to investigate anatomy in different contexts with the advantage of not having to apply the laborious traditional workflow (e.g., embedding, sectioning, staining) used for plant anatomical studies.

Because LATscan enabled direct visualization of the spatial variation of some (lignin and suber) cell wall components without any chemical treatment or staining of cell walls, it can be considered another tool to facilitate the integration of histology and chemical analysis of cell walls in plants. In addition to gross anatomy, this approach may be particularly useful for studying complex systems such as wounding experiments (e.g., grafting), plant-pathogen associations or parasitic plants-host interactions, which normally requires complementary chemical analysis (e.g., histochemistry, fluorescence microscopy) to identify tissues that are specific to the pathogen/parasitic plant or host plant, or to differentiate chemical compounds deposited in boundary layers of wounded plants (Rittinger *et al*., 1987; Rath *et al*., 2014; Navarro *et al*., 2019; Pellissari *et al*., 2022).

### The limitations of LATscan for plant anatomical studies

Most plant phenotyping techniques are destructive, and LAT is no different. Although LATscan allowed the adequate observation of most woody tissues, and in some cases improved the resolution of some structures (e.g., heartwood), in other cases the scanning was not efficient to perfectly illustrate the complexity of cell structures. For example, the resolution of vessels, parenchyma, sclerenchyma is adequate for studies at the tissue and cell levels, but g-fibers resulted in blurry cell patches in some cases. G-fibers have lignified outer secondary cell-wall layers, and a thick internal layer, the G-layer, that is formed mostly by cellulose (∼75% Mellerowicz & Gorshkova, 2012) that only later can become lignified (Guedes *et al*., 2017).

Because the G-layer is rich in non-structural polysaccharides such as pectin, the G-fiber cell wall forms a gel-like structure (Clair *et al*., 2008). This cell wall composition gives G-fibers a highly hydrophilic composition, and G-layers can easily shrink as moisture decreases (Clair *et al*., 2008; Guedes *et al*., 2017), which may generate the blurry aspect in LATscan.

Another limitation presents in the 3D reconstructions shown in Figure 5. In the axial direction of the blocks, striations are visible that make the longitudinal resolution not as sharp as the cross-sectional images. There are two reasons for this. First, the cross-sectional image at the laser ablation plane resolutions results in a resolution of 1 μm/pixel, whereas the sections are separated by 5 μm, thus leading to a reduced resolution in the longitudinal direction. Second, the UV-induced fluorescence is similar but not exactly the same in each section, which results in slight differences that manifest as these striations. The use of lasers with smaller pulse durations (we used a nanosecond pulsed laser in this study) in the picosecond and femtosecond range would help alleviate this issue with finer resolution in the longitudinal direction and more uniformity in the UV-induced fluorescence between sections.

Other disadvantage of LATscan compared to traditional anatomy is that, as a new technology, it may not be cost-effective compared to other traditional techniques (e.g., light microscopy). Nevertheless, as LATscan generate multiple data (2D, 3D, chemical composition information), a comparison in terms of cost would require including a full combination of techniques to acquire the same amount of data using traditional methods. In addition, 3D reconstructions are only possible using modern techniques (e.g., confocal laser scanning microscope, X-ray microtomography, magnetic resonance imaging) which might be equally not as cost-effective or cheaper. Here we optimized the application of LATscan for woody tissues which were obtained from stems with maximum diameter of 20 mm. However, such limitations regarding sample size are also real for nearly all other classical methods using microtomy and microscopy.

## Conclusions

The systematic approach described in this article opens new possibilities in the study of plant phenotyping. Laser ablation tomography (LATscan) is an innovative technique that allows for prompt, 2D and 3D image reconstructions of plant samples, offering new avenues to explore the complexity of plant morphology. This new technology will be particularly significant to filling in the gap of sample throughput that is not achieved by conventional microscopy techniques. Future plant anatomical research woody plants will benefit from the power and efficiency of LAT scans to illuminate the development, structure, chemical diversity, and 3D phenotypical information which remains strongly underused in plant sciences. Such characterizations will strength both basic and applied research, allowing in-depth investigations to be undertaken in areas ranging from plant anatomy to systems biology.

## Data availability

Data supporting the observations are largely presented in Supporting Information. Code and Dataset S2 are available at github.com/joycechery/LATScans. Movies are available at Zenodo repository (10.5281/zenodo.7289450). More detailed information, if necessary, will be provided on request.

## Acknowledgments

We would like to acknowledge the Arnold Arboretum of Harvard University for providing access to the living collections and financial support through a Sargent Award for Visiting Scholars (I.L.C.N). We thank Johanna M. Dela Cruz and the Cornell Institute of Biotechnology’s Imaging Facility for the support with the Spinning Disk Confocal Microscope (NIH 1S10OD010605) and Zeiss LSM 710 Confocal Microscope (NIH 1S10RR025502). This work was financially supported by Cornell University Lab Startup Funds (J.G.O). The work was also supported by University of Southern Maine startup funds provided by the Maine Economic Improvement Fund (A.L.).

## Author contributions

I.L.C.N. planned the research; I.L.C.N., B.H., A.L., J.B. and J.G.O. designed the research; I.L.C.N., B.H., A.L. and J.G.O. performed the research; I.L.C.N., B.H., A.L. and J.G.O. collected data and analyzed the data; I.L.C.N. and J.G.O. interpreted the results; I.L.C.N. wrote the paper with inputs from B.H., A.L. and J.G.O. All authors read and approved the final version of the manuscript.

## Supporting Information

**Table S1** Summary of traditional and modern techniques to study plant anatomy and cell biology.

**Table S2** Results of mean fluorescence ANOVA with post-hoc tests between all cell types.

**Fig. S1** Confocal laser scanning microscopy of samples of *Wisteria sinensis* to test infiltration time for safranin (a-c) and Calcofluor White (d-f). Emission wavelength and time of infiltration are given in each image.

**Fig. S2** LATscan of *Wisteria sinensis* stems compared by distinct staining methods. (A) Stained with Safrablau. (B) Stained with Calcofluor White. Stem diameter: 35 mm.

**Fig. S3**. Macroscopic image of *Wisteria sinensis* with heartwood. (A) Stem recently collected. (B) Fixed stem imaged with stereomicroscopy coupled with digital camera. (C) Detail of previous image. Stem diameter: 35 mm.

**Movie S1** Movie illustrating anastomoses and splitting between vascular cylinders in the compound stem of *Paullinia pinnata*.

**Movie S2** Movie illustrating the stem of woody vines in cross section and longitudinal radial section. The movie starts off in transverse view then reorients to a tangential view. Species name (from left to right): *Wisteria floribunda, Gnetum urens, Cocculus orbiculatus* and *Menispermum canadense*.

**Dataset S1** Results of particle analyzer for quantitative analysis of wood of *Urvillea chacoensis* (Sapindaceae). Counts are sorted from smaller to largest.

**Dataset S2** Measurements of emission wavelength of cell walls for different cell types from the stems of four different species of woody vines.

