## Supplemental for "Laser ablation tomography (LATscan) as a new tool for anatomical studies of woody plants"

**The following supporting information is available for this article**:

(Movies, Code & Dataset 2 are provided separately at: github.com/joycechery and at Zenodo repository (10.5281/zenodo.7289450):

**Table S1** Summary of traditional and modern techniques to study plant anatomy and cell biology.

**Table S2** Results of mean fluorescence ANOVA with post-hoc tests between all cell types.

**Fig. S1** Confocal laser scanning microscopy of samples of *Wisteria sinensis* to test infiltration time for safranin (a-c) and Calcofluor White (d-f). Emission wavelength and time of infiltration are given in each image.

**Fig. S2** LATscans of *Wisteria sinensis* stems compared by distinct staining methods. (A) Stained with Safrablau. (B) Stained with Calcofluor White. Stem diameter: 35 mm.

**Fig. S3**. Macroscopic image of *Wisteria sinensis* with heartwood. (A) Stem recently collected. (B) Fixed stem imaged with stereomicroscopy coupled with digital camera. (C) Detail of previous image. Stem diameter: 35 mm.

**Movie S1** Movie illustrating anastomoses and splitting between vascular cylinders in the compound stem of *Paullinia pinnata*.

**Movie S2** Movie illustrating the stem of woody vines in cross section and longitudinal radial section. The movie starts off in transverse view then reorients to a tangential view. Species name (from left to right): *Wisteria sp., Gnetum urens, Cocculus orbiculatus* and *Menispermum canadense.*

**Dataset S1** Results of particle analyzer for quantitative analysis of wood of *Urvillea chacoensis* (Sapindaceae). Counts are sorted from smaller to largest.

**Dataset S2** Measurements of emission wavelength of cell walls for different cell types from the stems of four different species of woody vines.

**Table S1** Review of traditional and modern techniques to study plant anatomy and cell biology with emphasis on woody plants.

| Approach | Required sample preparation | Imaging systems | Source | Additional references |
|---|---|---|---|---|
| Macroscopic analysis | 1. Wood block polishing | • Light microscopy (bright field)<br>• XyloPhone<br>• XyloScope | • Barbosa *et al.*, 2021<br>• Wiedenhoeft, 2020<br>• Hermanson *et al.*, 2019 | Luna-Márquez *et al.*, 2021; Nejapa *et al.*, 2021; Ravindran *et al.*, 2019; Ravindran & Wiedenhoeft, 2020 |
| | 2. Wood block trimming | • Light microscopy (epi-illumination) | • Lee *et al.*, 2022 | |
| Microscopic analysis | 1. Histological slides: fixation + embedding + sectioning + staining | • Light microscopy (bright field) | • Johansen, 1940; Ruzin, 1999 | Pace *et al.*, 2009; Crivellaro & Schweingruber, 2015; Angyalossy *et al.*, 2016; Chery *et al.*, 2020; Cunha Neto *et al.*, 2021 |
| | 2. Sanded surface: Wood block polishing | • Light microscopy/Reflected light/Fluorescence microscopy | • Kitin *et al.*, 2021 | |
| | 3. Sections: fixation + embedding + sectioning + coating | • Scannning electron microscopy | • Hatano *et al.*, 2022 | |
| Histochemical analysis | 1. Histological slides: fixation + sectioning + staining | • Light microscopy (bright field)<br>• Fluorescence microscopy | • Jensen, 1962; Ruzin, 1999; Demarco, 2017; Maceda & Terrazas, 2022 | Cunha Neto *et al.*, 2017; Pace *et al.*, 2019; Souza *et al.*, 2021 |
| | 2. Histological slides: sectioning + autofluorescence | • Confocal laser scanning microscopy | • Hutzler *et al.*, 1998 | Sando *et al.*, 2009; Nakazawa *et al.*, 2013 |
| Immunohistochemistry | 1. Sections: fixation + embedding + sectioning + antibody incubation | • Confocal laser scanning microscopy/ Fluorescence microscopy | • Hall *et al.*, 2013 | Verhertbruggen *et al.*, 2017 |
| | 2. Whole-mount fixation + permeabilization + antibody incubation | • Confocal laser scanning microscopy/ Fluorescence microscopy | • Sauer *et al.*, 2006; | Pasternak *et al.*, 2015 |
| 3D imaging | 1. Histological slides: Tissue isolation or sectioning (staining optional) | • Confocal laser scanning microscopy | • (Hepler & Gunning, 1998; Prunet *et al.*, 2016) | (Nakaba *et al.*, 2015) |

| | | | | |
|---|---|---|---|---|
| | 2. Whole sample: staining/phase-contrast (optional) | • X-ray microtomography<br>• Magnetic resonance microscopy<br>• Laser Ablation Tomography | • (Stuppy *et al.*, 2003)<br>• (Oven *et al.*, 2011)<br>• (Hall & Lanba, 2019) | Mayo *et al.*, 2010; Teixeira-Costa & Ceccantini, 2016; Mylo *et al.*, 2021; (Strock *et al.*, 2019; Martínez-Gómez *et al.*, 2022) |

**Table S2** Results of mean fluorescence ANOVA with post-hoc tests between all cell types.

**ANOVA**

| | **Df** | **Sum Sq** | **Mean Sq** | **F value** | ***P* value** |
|---|---|---|---|---|---|
| **Feature** | 4 | 110421 | 27605 | 40.06 | <2e-16 *** |
| **Residuals** | 250 | 172281 | 689 | | |

*P-value* indicate significance to the 99% level.

**Fig. S1** Confocal laser scanning microscopy of samples of *Wisteria sinensis* to test infiltration time for safranin (a-c) and Calcofluor White (d-f). Emission wavelength and time of infiltration are given in each image.

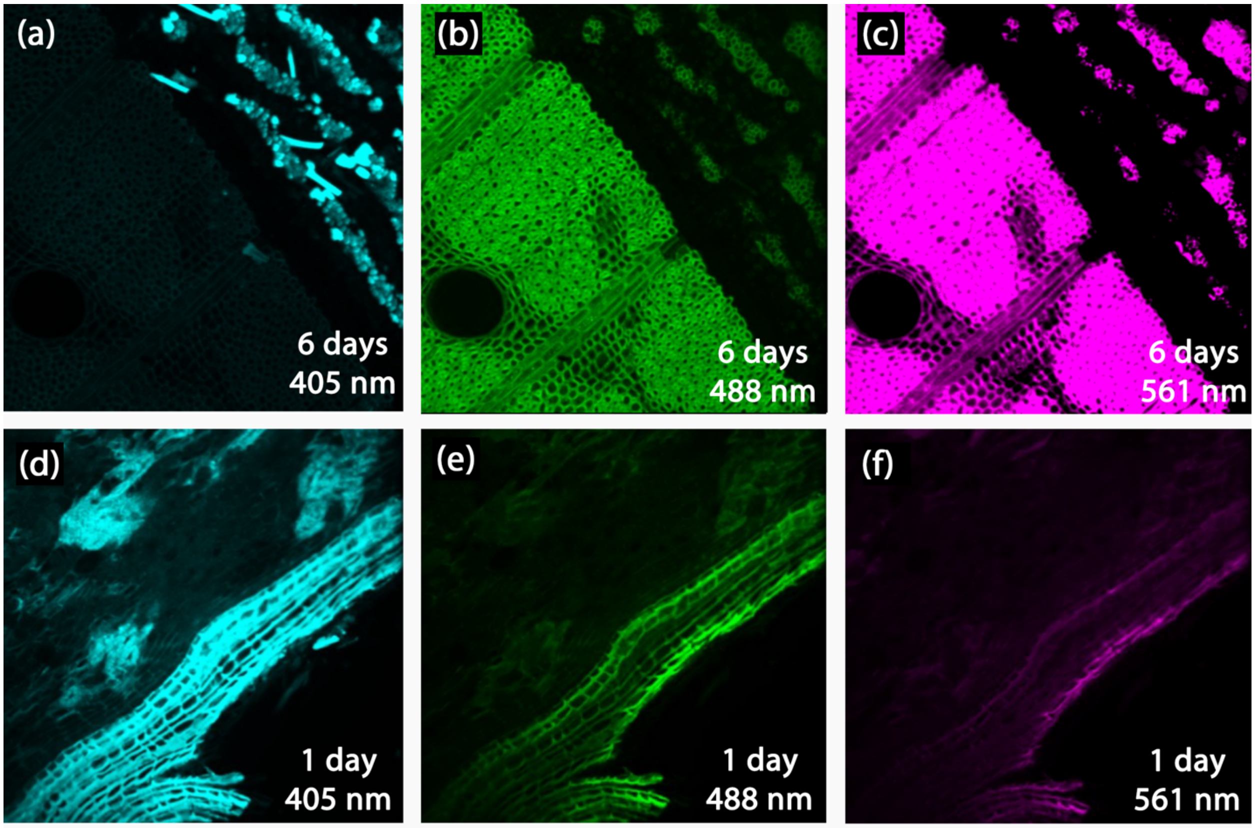

**Fig. S2** LATscans of *Wisteria sinensis* stems compared by distinct staining methods. (A) Stained with Safrablau. (B) Stained with Calcofluor White. Stem diameter: 35 mm.

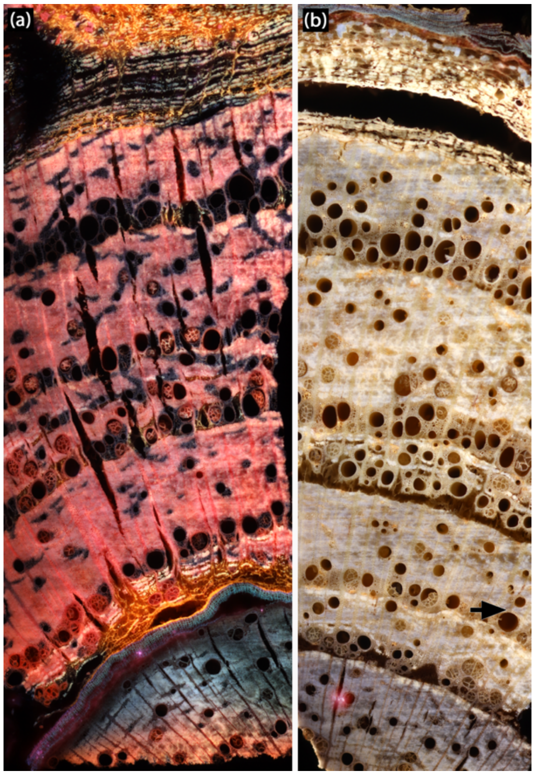


**Fig. S3**. Macroscopic image of *Wisteria sinensis* with heartwood. (A) Stem recently collected. (B) Fixed stem imaged with stereomicroscopy coupled with digital camera. (C) Detail of previous image. Stem diameter: 35 mm.

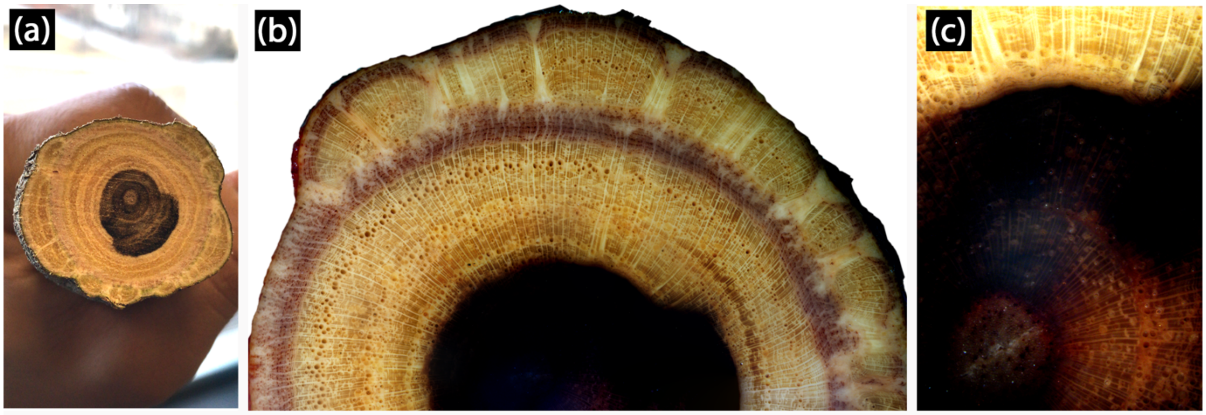


**Dataset S1** Results of particle analyzer for quantitative analysis of wood of *Urvillea chacoensis* (Sapindaceae). Counts are sorted from smaller to largest. See also figure (below) showing outlines that represent the vessels measured after segmentation of LATscan image. Counts 60, 15, 14, 36 and 34 were deleted as they were under the 50 microns threshold. Count 14 is not a vessel. Manual corrections were not performed.

| Count | Area | r^2 | r | diameter |
|---|---|---|---|---|
| 56 | 2251.068 | 716.5372 | 26.76821 | 53.53642 |
| 28 | 2333.06 | 742.6361 | 27.25135 | 54.5027 |
| 8 | 2407.599 | 766.3626 | 27.68325 | 55.36651 |
| 40 | 2611.338 | 831.2147 | 28.83079 | 57.66159 |
| 57 | 2616.307 | 832.7964 | 28.85821 | 57.71642 |
| 11 | 3105.778 | 988.5998 | 31.44201 | 62.88402 |
| 29 | 3195.224 | 1017.071 | 31.89156 | 63.78311 |
| 39 | 3321.94 | 1057.406 | 32.51779 | 65.03557 |
| 50 | 3637.487 | 1157.848 | 34.02717 | 68.05433 |
| 9 | 3724.449 | 1185.529 | 34.43151 | 68.86302 |
| 31 | 4703.39 | 1497.136 | 38.69284 | 77.38567 |
| 17 | 5289.761 | 1683.783 | 41.03393 | 82.06786 |
| 49 | 5823.954 | 1853.822 | 43.05603 | 86.11207 |
| 30 | 5933.278 | 1888.621 | 43.45827 | 86.91654 |
| 24 | 6186.709 | 1969.291 | 44.37669 | 88.75338 |
| 47 | 6420.264 | 2043.634 | 45.20656 | 90.41313 |
| 26 | 7319.697 | 2329.932 | 48.26937 | 96.53874 |
| 58 | 7597.975 | 2418.511 | 49.17835 | 98.35671 |
| 44 | 7707.298 | 2453.309 | 49.53089 | 99.06178 |
| 32 | 8537.162 | 2717.463 | 52.12929 | 104.2586 |
| 51 | 8758.293 | 2787.851 | 52.80011 | 105.6002 |
| 48 | 9734.75 | 3098.667 | 55.66567 | 111.3313 |
| 52 | 9854.012 | 3136.629 | 56.00562 | 112.0112 |
| 33 | 10432.93 | 3320.904 | 57.62729 | 115.2546 |
| 13 | 10721.15 | 3412.646 | 58.41786 | 116.8357 |
| 45 | 11941.09 | 3800.968 | 61.65199 | 123.304 |
| 18 | 12204.46 | 3884.802 | 62.32818 | 124.6564 |
| 38 | 13111.35 | 4173.473 | 64.60242 | 129.2048 |
| 25 | 13575.98 | 4321.367 | 65.73711 | 131.4742 |
| 21 | 14212.04 | 4523.833 | 67.25944 | 134.5189 |
| 53 | 14810.83 | 4714.435 | 68.66174 | 137.3235 |
| 1 | 15238.19 | 4850.466 | 69.64529 | 139.2906 |
| 22 | 15722.69 | 5004.687 | 70.74381 | 141.4876 |
| 5 | 16028.3 | 5101.966 | 71.42805 | 142.8561 |
| 46 | 16030.78 | 5102.756 | 71.43358 | 142.8672 |
| 41 | 16475.53 | 5244.324 | 72.41771 | 144.8354 |
| 42 | 17437.08 | 5550.394 | 74.50097 | 149.0019 |
| 43 | 17750.14 | 5650.045 | 75.16678 | 150.3336 |
| 54 | 19069.48 | 6070.002 | 77.91022 | 155.8204 |
| 12 | 19357.69 | 6161.744 | 78.49678 | 156.9936 |
| 35 | 20393.78 | 6491.541 | 80.5701 | 161.1402 |
| 23 | 21516.83 | 6849.019 | 82.7588 | 165.5176 |
| 20 | 21593.85 | 6873.536 | 82.90679 | 165.8136 |
| 27 | 21807.53 | 6941.552 | 83.31598 | 166.632 |
| 59 | 22413.78 | 7134.527 | 84.46613 | 168.9323 |

| | | | | |
|---|---|---|---|---|
| 19 | 23223.76 | 7392.353 | 85.9788 | 171.9576 |
| 3 | 24061.08 | 7658.88 | 87.51503 | 175.0301 |
| 55 | 27015.3 | 8599.236 | 92.73207 | 185.4641 |
| 7 | 31606.88 | 10060.78 | 100.3034 | 200.6069 |
| 10 | 33105.11 | 10537.68 | 102.6532 | 205.3064 |
| 37 | 34285.3 | 10913.35 | 104.467 | 208.934 |
| 6 | 34528.79 | 10990.86 | 104.8373 | 209.6746 |
| 4 | 35592.21 | 11329.35 | 106.4394 | 212.8789 |
| 2 | 35790.98 | 11392.62 | 106.7362 | 213.4725 |
| 16 | 41791.35 | 13302.6 | 115.3369 | 230.6738 |

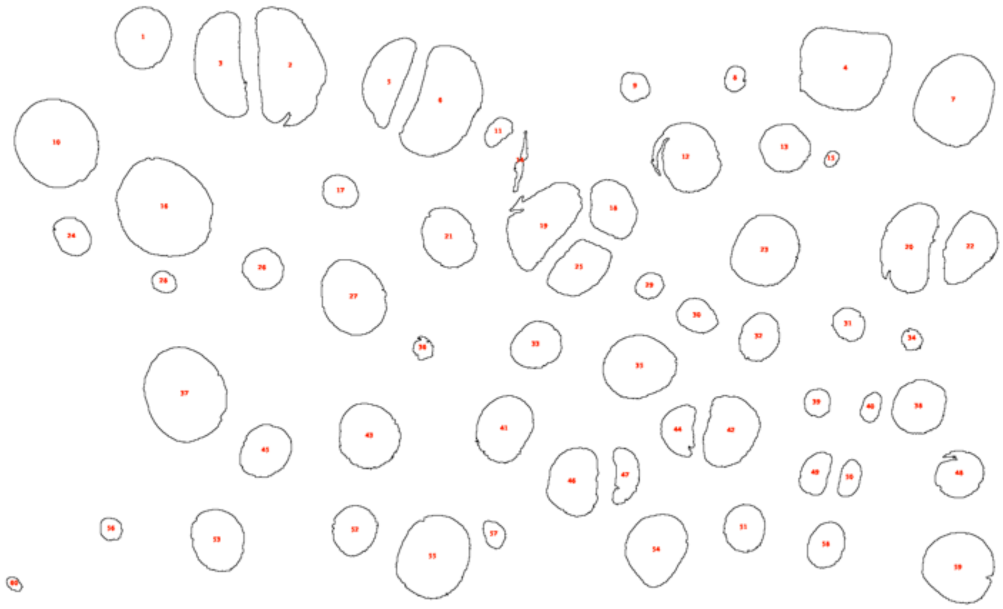
